# Conserved circadian membrane rhythms arise from divergent cellular mechanisms in pacemaker neurons of mice and *Drosophila*

**DOI:** 10.64898/2026.02.04.703843

**Authors:** Edgar Buhl, Christian Nathan, Hugh D Piggins, Krasimira Tsaneva-Atanasova, Mino DC Belle, James JL Hodge

**Affiliations:** School of Physiology, Pharmacology and Neuroscience, University of Bristol, UK; Centre for Biological Timing, Division of Neuroscience, School of Biological Sciences, Faculty of Biology, Medicine and Health, University of Manchester, UK; Department of Mathematics and Statistics and Living Systems Institute, University of Exeter, UK

**Keywords:** circadian rhythms, membrane excitability, ion channels, neurophysiology, post-inhibitory rebound

## Abstract

Circadian clocks align physiology and behaviour with daily environmental cycles, requiring neuronal networks that reliably encode and transmit time-of-day information. A fundamental yet unresolved question is how membrane properties support this function and whether conserved principles arise across evolutionarily distant organisms. We compared the intrinsic electrophysiological properties of identified circadian neurons from a nocturnal mammal (mouse, *Mus musculus* SCN VIP neurons) and a diurnal insect (fly, *Drosophila melanogaster* l-LNv PDF neurons) across the 24-h light-dark cycle. Despite ∼700 million years of evolutionary divergence and opposite temporal niches, we found neurons from both species exhibited similar rhythmic patterns in key parameters for encoding time-of-day, including resting membrane potential, spontaneous firing rate and cell capacitance, indicating convergent strategies for generating daily variation in excitability despite dramatically different network scales (∼2200 VIPs vs. 8 l-LNvs). These shared outputs arose through distinct mechanisms: flies exhibited higher input resistance, greater excitability and relatively larger A-type potassium currents, while mice displayed larger sustained outward currents and post-inhibitory rebound excitation, absent in flies. Rheobase troughs occurred during the inactive phase in both species, while other parameters did not cycle. These differences likely shape how each neuronal type integrates salient inputs, consistent with their respective roles in processing sustained daytime light (flies) versus nocturnal cues (mice). Our findings reveal conserved functional goals but divergent electrophysiological strategies in circadian neurons, reflecting evolutionary adaptation to species-specific environmental demands. Such neurophysiological diversity highlights how evolution shapes cellular mechanisms to meet the requirements of temporal niche while preserving robust circadian timekeeping.

## Introduction

Circadian rhythms are a universal feature of life, from unicellular organisms to humans. They operate at molecular, cellular and electrophysiological levels, and reflect the evolutionary adaptation of organisms to predictable daily and seasonal environmental changes caused by the Earth’s rotation and orbit around the Sun. To cope with these changes, organisms rely on circadian clocks - intrinsic timekeeping systems that regulate behaviour, physiology and metabolism - ensuring that these processes occur at optimal times of the day, enhancing an organism’s fitness (Dunlap *et al*., 2004; Allada and Chung, 2010). The circadian system is typically organised into three interacting components: inputs that synchronise the clock to external cues, a central oscillator, and outputs that regulate rhythmic physiology and behaviour. At the core of circadian timekeeping lies the molecular clock, a cell-autonomous transcription-translation feedback loop (TTFL) in which rhythmically expressed clock genes regulate the circadian transcription of downstream targets, including ion channels, receptors and transporters (Hegazi *et al*., 2019). This mechanism, well-conserved across evolution, has been extensively studied in the fruit fly *Drosophila melanogaster* and later shown to operate similarly in mice and humans (Dunlap, 1999; Panda *et al*., 2002; Takahashi, 2017).

Mice and flies represent two evolutionarily distinct model systems - mammals and invertebrates - that nevertheless share a highly conserved molecular circadian clock. They also occupy different temporal niches: mice are nocturnal and avoid light (Tamayo *et al*., 2023), whereas *Drosophila* are diurnal and actively seek it (Parsons, 1975) (Figure 1). Consequently, although light serves as a primary zeitgeber in both species, it is encountered under very different temporal and behavioural conditions, with strong effects on neuronal excitability and network dynamics. Day-active flies experience light predominantly during their active phase, while nocturnal mice primarily encounter it outside their active phase, requiring circadian circuits to adjust appropriately to these species-specific environmental demands. Despite these differences, both species have been instrumental in advancing our understanding of circadian clocks. A major distinction lies in the overall organisation of their clock circuitry: the mammalian suprachiasmatic nucleus (SCN) contains ∼20,000 heterogenous neurons in a densely packed anatomically defined brain region, comprising core and shell subregions (Antle and Silver, 2005), while the *Drosophila* brain harbours only ∼240 individually identifiable clock neurons distributed in a dispersed neural network with discrete dorsal and lateral clusters (Helfrich-Förster and Reinhard, 2025) (Figure 1). These neuronal populations differ in their anatomical positions, projection patterns, and neurochemical identities, yet both clock networks share common operating principles of synchronisation, excitability, and time-of-day signalling.

**Figure 1:**
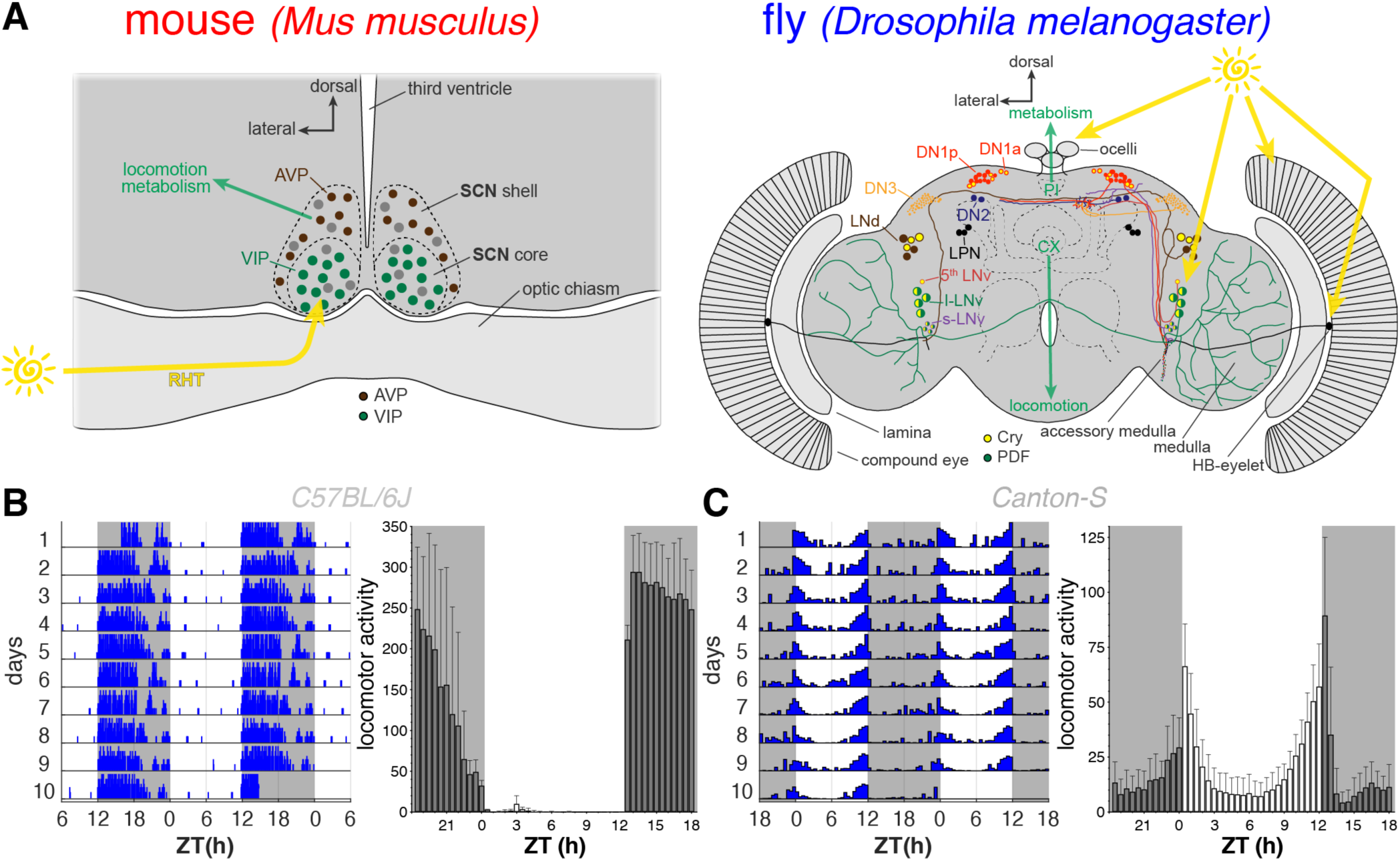
Mouse and fly circadian clock circuit and light-dark activity patterns. (**A**) Simplified schematic of the mammalian and *Drosophila* circadian clocks. The mammalian master clock resides in the suprachiasmatic nucleus (SCN, left), comprising ∼20,000 heterogenous neurons organised into a ventral core region containing VIP neurons and a dorsal shell region primarily containing AVP neurons. Core and shell neurons are synchronised via multiple intercellular communication mechanisms, enabling coherent outputs to peripheral clocks. The *Drosophila* clock (right) consists of 240 individually identifiable neurons arranged into a distributed network of distinct groups with specific functions, symmetrically distributed in each brain hemisphere: dorsal lateral neurons (LNd); large and small PDF-expressing ventral lateral neurons (LNv); a single PDF-negative ventral lateral neuron (5th LNv); lateral posterior neurons (LPN); dorsal neurons 1 (DN1a and DN1p); dorsal neurons 2 (DN2); and dorsal neurons 3 (DN3). In mice, light input from the eyes to the SCN is primarily glutamatergic via the retinohypothalamic tract (RHT), mainly targeting ventral core VIP neurons. In flies, light is detected by the cell-autonomous blue-light photoreceptor cryptochrome (Cry), extraretinal photoreceptors in the HB-eyelet, and the compound eyes and ocelli, all converging on the LNvs. Yellow and green arrows indicate major input and output pathways, respectively. AVP, arginine vasopressin; Cry, cryptochrome; PDF, pigment dispersing factor; VIP, vasoactive intestinal peptide. (**B**, **C**) Double-plotted actograms of a representative wild-type mouse (*C57BL/6J*, **B**) wheel-running activity and a wild-type fly (*Canton-S*, **C**) locomotor activity recorded over ten days in a 12 h:12 h light–dark cycle (white, lights on; grey, lights off; left panels), with corresponding histograms showing 10-day average activity from 3 mice and 20 flies, respectively, illustrating their characteristic nocturnal (mouse) and diurnal–crepuscular (fly) activity patterns (right panels). Bars, mean; error bars, SD.

While individual neurons can function as autonomous circadian clocks, rhythmic behaviour arises from networks of interacting neurons that communicate through electrical activity, synaptic, and paracrine transmission. In both mammals and *Drosophila*, the molecular clock drives daily changes in neuronal excitability by regulating intracellular signalling cascades, which in turn affect ion channel activity and neurotransmitter release, ultimately generating time-of-day signals (Harvey *et al*., 2020). These are expressed as rhythms in the passive and active membrane properties of clock neurons, including resting membrane potential (RMP), input resistance (R_in_), neuronal excitability, and ionic currents (Allen *et al*., ^2^01^7^). Beyond these basic measures, properties such as spike-frequency adaptation, post-inhibitory rebound (PIR), and voltage-gated current amplitudes are critical determinants of how clock neurons encode time-of-day signals and process synaptic inputs (Harvey *et al*., 2020). Clock neurons are typically more depolarised during the day than at night, with spontaneous firing rate (SFR) providing a direct readout of circadian phase (Belle and Diekman, 2018). Together, these intrinsic electrophysiological properties shape synaptic integration and signal processing, determining how circadian signals are transmitted within and beyond the clock network. In doing so, electrical membrane rhythms synchronise individual clock neurons, broadcast timing signals across the brain and body, and provide feedback to the molecular oscillator itself (Mizrak *et al*., 2012). Strikingly, silencing electrical activity disrupts molecular oscillations and abolishes behavioural rhythms (Nitabach *et al*., 2002; Depetris-Chauvin *et al*., 2011), pointing to two interdependent circadian pacemakers - the intracellular molecular and the membrane electrical oscillator - functioning as the two hands of the circadian clock. While the molecular hand is private to each cell, the membrane hand is public and broadcasts circadian information.

Despite differences in organisation, functionally, the similarities between mammalian and fly clock neurons are striking. SCN neurons synchronise through GABAergic synapses, peptidergic signals such as vasoactive intestinal peptide (VIP) and arginine vasopressin (AVP), and gap-junctional coupling (Honma, 2018), whereas *Drosophila* clock neurons also communicate via classical neurotransmitters and the neuropeptide pigment dispersing factor (PDF), a major synchronising signal through its widespread receptor expression (Helfrich-Förster and Reinhard, 2025). Innexins, the invertebrate homologues of vertebrate connexins, mediate electrical gap junction coupling, with Innexin2 expressed in PDF-positive ventral lateral neurons (LNv) modulating free-running period length (Ramakrishnan and Sheeba, 2021), while blocking gap junctions reduces firing frequency (Cao and Nitabach, 2008). Together, these parallels highlight the conserved role of electrical and chemical coupling in circadian circuit function across phyla.

To date, several studies have reported diurnal variation in neurophysiological parameters of clock neurons such as RMP, SFR and input resistance in both mice (Kuhlman and McMahon, 2004; Belle *et al*., 2009; Flourakis *et al*., 2015; Hermanstyne *et al*., 2016; Wegner *et al*., 2017; Patton *et al*., 2020; Hermanstyne *et al*., 2023; Yang *et al*., 2023) and flies (Park and Griffith, 2006; Cao and Nitabach, 2008; Sheeba *et al*., 2008a; Sheeba *et al*., 2008b; Flourakis *et al*., 2015; Buhl *et al*., 2016; Buhl *et al*., 2019; Curran *et al*., 2019; Buhl *et al*., 2022; Tang *et al*., 2022). However, the findings are not always consistent, with different laboratories sometimes reporting divergent results. These discrepancies likely reflect differences in experimental conditions, recording time points, recording techniques, and analysis protocols, which complicate direct comparison within and particularly between species.

To enable robust cross-species comparisons, we systematically compare the neurophysiological properties of mouse VIP neurons in the ventral SCN core (approx. 2200 neurons (Abrahamson and Moore, 2001)) and *Drosophila* PDF-positive large LNvs (8 neurons (Helfrich-Förster, 1995)) across the 24 h light-dark cycle under matched conditions and analysis – to provide a level of consistency typically unavailable in previous studies. VIP and l-LNv neurons are well-characterised clock neurons that play central roles in light-input pathways (Parisky *et al*., 2008; Sheeba *et al*., 2008b; Buhl *et al*., 2016; Schlichting *et al*., 2016; Jones *et al*., 2018; Collins *et al*., 2020; Paul *et al*., 2020). Notably, VIP and PDF signal via homologous G-protein–coupled receptors (VPAC2 and PDFR respectively) that elevate intracellular cAMP levels, acting as key synchronising signals with phase-delaying effects, thereby fulfilling analogous roles in circadian network organisation (Helfrich-Förster and Reinhard, 2025). By comprehensively measuring a broad set of active and passive membrane properties, including excitability, ionic currents, input resistance, and membrane dynamics, across the day and night, we establish a framework to test the hypothesis that, while core biophysical principles of circadian membrane function are conserved, individual clock neurons display species-specific electrophysiological adaptations shaped by their distinct temporal niches, photoreceptive pathways, and circuit architectures.

## Materials and methods

We kept recording solutions, protocols, equipment and procedures as similar as possible between mouse and fly preparations, in order to facilitate direct comparisons and to ensure that observed differences and similarities in clock neuron physiology reflect biological rather than methodological variation. Nonetheless, some adaptations were necessary to account for species-specific requirements in tissue preparation and handling, which are detailed below.

### Animals

#### Mouse

All animal procedures were conducted in accordance with the UK Animals (Scientific Procedures) Act of 1986 and approved by the University of Manchester Ethics Committee and the Home Office (project license PP5674327). Experimental animals (VIP-eYFP, in which VIP-expressing neurons are genetically labelled with eYFP) were generated by crossing VIP-ires-Cre mice (JAX #010908; (Taniguchi *et al*., 2011)) with mice carrying a Cre-dependent eYFP reporter allele (JAX #006148; R26R-eYFP, B6.129X1-Gt(ROSA)26Sor^tm1(eYFP)Cos^/J). Parent strains were backcrossed for 5-6 generations onto a C57BL/6J background, which also served as the wild-type behavioural control, to reduce genetic variability. Animals were bred and maintained in the University of Manchester animal facility. Adult mice (*Mus musculus*, 3-5 months old) were group-housed in mixed-sex cohorts in light-tight and soundproof cabinets (Tecniplast LTD, UK) under a 12 h:12 h light-dark (LD) cycle at 22°C, with food and water available *ad libitum*.

#### Fly

Flies (*Drosophila melanogaster*) of the *PDF-RFP strain* (Ruben *et al*., 2012) and *Canton-S w+* (behavioural wild-type control) were reared under a 12 h:12 h LD cycle on standard medium (0.7% agar, 1.0% soya flour, 8.0% polenta/maize, 1.8% yeast, 8.0% malt extract, 4.0% molasses, 0.8% propionic acid, 2.3% nipagen) at 25°C. Adult flies were collected 2-5 days post-eclosion for experiments.

Zeitgeber time (ZT) 0 was defined as lights on and ZT12 as lights off. Behavioural experiments were conducted with 3 male mice and 20 male flies. Electrophysiology experiments were performed using 12 female and 11 male mice, and 48 female and 50 male flies.

### Locomotor behaviour

#### Mouse

Wheel-running activity was recorded from individually housed male wild-type C57BL/6J mice (3-4 months old; purchased from Harlan, Blackthorn, UK) using cages equipped with stainless-steel running wheels (16 cm diameter) and grid floors to allow collection of food and waste without disturbing the home-cage environment (North Kent Plastics, Kent, UK). Activity was continuously monitored and binned in 10-minute intervals using the Chronobiology Kit (Stanford Software Systems, Santa Cruz, CA). Mice were entrained and maintained under a 12 h:12 h LD cycle (lights-on: 250 lux) at 23°C and 40% relative humidity for ten days.

#### Fly

Circadian locomotor activity was recorded using the *Drosophila* Activity Monitor system (DAM2, Trikinetics Inc., Waltham, MA, USA). Individual entrained male flies (*Canton-S w+*) were placed in DAM tubes containing a small amount of standard food. The monitors were housed in a light- and temperature-controlled incubator (Percival Scientific Inc., Perry, IA, USA), and fly activity was recorded for ten days under 12 h:12 h LD cycles at 25°C and 70% relative humidity.

### Brain and tissue preparation for electrophysiological recordings

#### Mouse

Mice were sedated with isoflurane (Abbott Laboratories) and euthanised by cervical dislocation either during the light phase or, for dark-phase experiments, under infrared illumination. Brains were rapidly removed, mounted on a metal stage, and sectioned into 250 µm coronal slices encompassing the mid-suprachiasmatic nucleus (SCN) across the rostro-caudal axis using a vibrating microtome (Leica VT1200, Leica Microsystems Ltd, Germany). Slices were cut in ice-cold (4°C) sucrose-based incubation solution containing (in mM): 95 NaCl, 1.8 KCl, 0.5 CaCl_2_, 7 MgSO_4_, 1.2 KH_2_PO_4_, 26 NaHCO_3_, 15 glucose, 50 sucrose, 0.005 mg/L Phenol Red, continuously oxygenated with 95% O_2_, 5% CO_2_, pH 7.4. After slicing, the tissue was transferred to a holding chamber with the same oxygenated solution at room temperature (20-23°C) for ≥20 min, then transferred to recording artificial cerebrospinal fluid (aCSF). The recording aCSF composition was identical to the incubation solution except for the following (mM): 127 NaCl, 2.4 CaCl_2_, 1.3 MgSO_4_, 0 sucrose. Measured osmolarity for both incubation and recording solutions was 300-310 mOsmol/kg. Slices were allowed to recover in aCSF for ≥90 min before recordings.

#### Fly

Adult flies were briefly anaesthetised with CO_2_, decapitated, and brains rapidly removed. Whole brains were dissected and cleaned in extracellular saline at room temperature containing (in mM): 101 NaCl, 3 KCl, 1 CaCl_2_, 4 MgCl_2_, 1.25 NaH_2_PO_4_, 20.7 NaHCO_3_, 5 D-glucose, pH 7.2. Photoreceptors, lamina, air sacs, trachea, and the ganglion sheath were carefully removed using sharp forceps. Enzymatic treatment was avoided to preserve the extracellular matrix and maintain the integrity of membrane proteins, including ion channels and receptors. A small incision was made at the central brain-medulla border above the l-LNv neurons to facilitate electrode access. For dark phase recordings (ZT13-24), flies and preparations were handled under red safe-light illumination, used both for ambient lighting and dissection microscope filters.

### Whole-cell patch clamp recordings

Targeted whole-cell current- and voltage-clamp recordings were obtained from VIP-eYFP positive neurons in acute mouse SCN brain slices and from large ventrolateral clock neurons (l-LNvs) in *Drosophila* brains. Recordings were performed across the circadian cycle (ZT1-24), encompassing both the day (ZT1-12) and night (ZT13-24). Experiments were conducted using a MultiClamp 700B amplifier (Molecular Devices, CA, USA) and digitised with a DigiData 1550B interface. Data were acquired with pClamp and MultiClamp Commander software at 20 kHz sampling and 10 kHz Bessel filtering. Unless otherwise stated, chemicals were obtained from Sigma (Poole, UK) or Avantor (Lutterworth, UK). The start time of each recording was used to assign the corresponding ZT. The procedures for tissue preparation and neuron identification differed slightly between species and are described in detail below. Only neurons that were capable of generating action potentials in response to depolarising current injection were included in the analysis, regardless of whether they exhibited spontaneous firing of action potentials at rest.

#### Mouse

SCN VIP-eYFP neurons were identified by fluorescence and visualised using an upright microscope (Olympus BX51 WI, Japan) equipped with a 40× water-immersion objective (Olympus, LUMPlanFLN, 40x/0.8), infrared differential interference contrast (IR-DIC) optics, fluorescence light source (pE4000, Cool LED, UK) and a cooled sensitive camera (Retiga Electro, Teledyne Photometrics). Slices were placed in the recording chamber, secured with a harp, and continuously perfused with oxygenated aCSF at ∼2.5 ml/min at room temperature.

Patch pipettes (7-8 MΩ) were pulled from thick-walled borosilicate glass (Harvard Apparatus, USA) using a two-stage puller (PC-100, Narishige, Japan). Electrodes were filled with intracellular solution containing (in mM): 120 K-gluconate, 20 KCl, 2 MgCl_2_, 2 K_2_-ATP, 0.5 Na-GTP, 10 HEPES, and 0.5 EGTA (pH 7.3 with KOH; 295-300 mOsmol/kg). Series resistance (15-20 MΩ) was compensated using bridge balance in current-clamp recordings, while cell capacitance was left uncompensated. In voltage-clamp, whole-cell capacitance (1.4-7.5 pF at 20-90 MΩ) and series resistance (50-75% at 5 kHz) were compensated. Recordings were included in the analysis only when access resistance was < 50 MΩ, leak current in response to a -40 mV pulse was < 50 pA, and the resting potential remained stable. In total, recordings were obtained from 302 ventral SCN VIP neurons (up to 15 cells per brain slice).

#### Fly

Large LNv neurons were visualised using an upright microscope (Examiner Z1, Zeiss, Germany) equipped with a 63× water-immersion objective, far-red illumination, and a cooled camera (optiMOS, QImaging, UK) and identified by RFP expression, size and position. Dissected brains were placed ventral side up in the recording chamber, secured with a custom-made brain harp, and maintained in a static bath of extracellular saline at room temperature (20-22 °C).

Patch pipettes (8-14 MΩ) were pulled from thick-walled borosilicate glass (World Precision Instruments, FL, USA) using a two-stage puller (PC-100, Narishige, Japan) and filled with intracellular solution containing (in mM): 102 K-gluconate, 17 NaCl, 0.94 EGTA, 8.5 HEPES, 0.085 CaCl_2_, 1.7 MgCl_2_ or 4 Mg·ATP and 0.5 Na·GTP, pH 7.2. Series resistance (15-25 MΩ) and cell capacitance (6 pF) were compensated using bridge balance in current-clamp recordings. In voltage-clamp, whole-cell capacitance (1.2-4 pF at 20-150 MΩ) and series resistance (50-75% at 5 kHz) were compensated. Recordings were included in the analysis only when access resistance was < 60 MΩ, leak current in response to a -40 mV pulse was < 60 pA, and the resting potential remained stable. In total, recordings were obtained from 288 LNv neurons, with a maximum of four cells recorded per brain.

#### Pharmacology

For voltage-clamp experiments measuring A-type potassium currents, tetrodotoxin (TTX) was applied to block fast voltage-gated sodium channels that would otherwise mask the potassium current. TTX (Tocris or Hello Bio, Bristol, UK) stock solutions were prepared in aCSF for mouse experiments and in external saline for fly experiments. For mouse recordings, TTX was diluted to a final concentration of 1 µM in aCSF immediately before bath application and delivered via continuous perfusion. For fly recordings, a 300 µM stock solution was added directly to the ∼1 ml static recording chamber to achieve a final concentration of 3-4 µM.

### Data analysis

Recordings were obtained from both male and female mice and flies; as no sex-dependent differences were observed in any parameter, data were pooled for statistical analysis. All electrophysiological analyses were performed in ClampFit (Molecular Devices) using identical procedures for both species to ensure comparability. The liquid junction potential was measured and calculated as 13 mV for both systems and both corrected and uncorrected membrane voltages in current-clamp recordings and command voltages in voltage-clamp experiments are reported. Spontaneous activity was measured in current-clamp mode with no holding current. During break-in and between voltage-clamp protocols, cells were held at -63 mV (fly) or -73 mV (mouse). Break-in was achieved using the membrane test tool in pClamp, and after stabilising the recording for 1 min, access resistance and leak currents were measured to assess recording quality.

#### Current-clamp

Average resting membrane potential (RMP) and spontaneous firing rate (SFR) were calculated over a 1 min recording period. The membrane time constant (τ) was obtained from the voltage response to a brief hyperpolarising current pulse (-100 pA, 20 ms). Five traces were averaged and fitted with up to three-exponential function using the Levenberg-Marquardt (LM) method from the end of the pulse until return to baseline (500-1000 ms). Goodness of fit was visually inspected, and τ was taken from the dominant (slowest) exponential component. Membrane input resistance (R_in_) was calculated from the steady-state voltage deflection in response to hyperpolarising current steps (-30 to -10 pA, 1 s) using Ohm’s law, and cell capacitance was subsequently estimated as τ/R_in_. The hyperpolarisation-activated current (I_h_) was quantified by applying a -30 pA current step (1 s). I_h_ amplitude was defined as the difference between the negative peak voltage within the first 300 ms of the step and the mean voltage over the final 100 ms. Neuronal excitability (f at +30 pA) was assessed by injecting 1 s depolarising current pulses of increasing amplitude up to +30 pA from rest and manually counting the resulting action potentials. Neuronal gain was determined as the slope of the linear portion of the resulting frequency-current (f-I) relationship.

Rheobase and responses to sinusoidal current stimulation were assessed with cells held at -73 mV (-60 mV LJP uncorrected). Rheobase was defined as the minimal current required to elicit the first action potential in response to a 50 ms depolarising pulse, increased in 1 pA increments. For sinusoidal stimulation, trains of 10 cycles were delivered at increasing frequencies (1, 2, 5, and 10 Hz) within a 30 s recording. These trains were repeated at increasing current amplitudes (+5, +15, +25 pA), and resulting spikes were manually counted. For analysis, a stimulation level was selected such that the highest frequency (10 Hz) elicited approximately five spikes. Spikes were then counted for each cycle across all trains at this current level. For cross-comparison, spike numbers were normalised to ten spikes at 1 Hz (one spike per cycle), and spike counts at higher frequencies were divided by the number of 1 Hz spikes to calculate the fraction of spikes per cycle.

#### Voltage-clamp

Current densities were calculated by dividing the recorded currents by the cell capacitance, measured in current-clamp mode at the start of each experiment. A-type potassium currents (I_A_) were measured in the presence of TTX using two voltage-clamp protocols. From a holding potential of -70 mV (-57 mV uncorrected), a 50 ms pre-step to -110 mV (-97 mV uncorrected) was applied, followed by a series of 250 ms voltage steps from -100 to +60 mV (-87 to +73 mV uncorrected) to evoke the total potassium current (I_total_). The same protocol was then repeated from a holding potential of +30 mV (+43 mV uncorrected) to isolate potassium currents lacking the A-type component (I_non-A_). The A-type current (I_A_) was obtained by digital subtraction of I_non-A_ from I_total_. Peak I_A_ amplitude was measured at the +60 mV (+73 mV uncorrected) step, and fast and slow time constants (τ_fast_ and τ_slow_) were determined by fitting a double-exponential function from the peak to 100 ms later using the LM model, with all proportions and time constants constrained to be positive. For sustained currents (I_sus_), traces were corrected for leak. Leak currents were measured by holding cells at -50 mV and applying 500 ms voltage steps from -60 to -140 mV (-47 to -127 mV uncorrected), with resistance calculated from the resulting I-V relationship using Ohm’s law. For I_sus_ measurements, cells were held at −70 mV (-57 mV uncorrected) and 500 ms voltage steps were applied from -100 to +60 mV (-87 to +73 mV uncorrected), with current measured over the final 100 ms of each step.

### Statistics

Values are reported as mean with 95% confidence intervals. Sample sizes are indicated in the text, figure legends, and tables. Statistical significance was considered at *p* < 0.05. All statistical analyses were performed in Prism (GraphPad Software Inc., CA, USA), and figures were arranged in Illustrator (Adobe Systems Inc., CA, USA). For presentation, voltage-clamp data were filtered at 3 kHz. Curve fitting was performed by comparing a sine-wave model with a non-zero baseline and a fixed 24 h period (representing a diurnal effect) against a linear model (representing no day/night difference). Model selection was based on Akaike’s Information Criterion (AIC). From the sine-wave model, amplitude, MESOR (Midline Estimating Statistic Of Rhythm), and phase shift (used to calculate acrophase) were obtained. The coefficient of determination (R^2^) is presented as a measure of goodness of fit. The original 24 h data were concatenated twice in order to improve fitting robustness; however, only a single 24 h cycle is presented in the figures. Simple linear regression was used to examine relationships between firing rate and membrane potential. Slopes were tested for deviation from zero and for differences between groups. For sinusoidal stimulation, two-way ANOVA was applied to assess differences between mouse and fly responses. For voltage-clamp comparisons across conditions (midday vs. midnight; mouse vs. fly), datasets were first assessed for compliance with parametric assumptions using the Shapiro-Wilk test (α = 0.05). Where assumptions were met, analyses were performed using ordinary one-way ANOVA; otherwise, the non-parametric Kruskal-Wallis test was applied. *Post hoc* comparisons were conducted using Sidak’s test (parametric) or Dunn’s test (non-parametric), as appropriate.

## Results

### Conserved daily rhythms in membrane potential and firing

To understand how conserved and divergent mechanisms shape the electrophysiological properties of circadian neurons, we compared clock neurons that occupy analogous positions in the light-input pathway of two evolutionarily distinct species: the mammalian mouse and the invertebrate fruit fly *Drosophila* (Figure 1A). Mice are nocturnal, showing most of their active behaviour during the dark phase (Figure 1B), whereas flies are diurnal, being behaviourally more active during the light phase (Figure 1C). These distinct temporal niches provide a natural context for examining potential species-specific adaptations in intrinsic neuronal properties.

To systematically compare their intrinsic electrophysiological properties, we performed whole-cell current- and voltage-clamp recordings from fluorescently labelled VIP (*VIP-eYFP*) and l-LNv (*PDF-RFP*) neurons across the 24 h light-dark (LD) cycle, binning data into 1 h zeitgeber time (ZT) intervals (Figure 2A). Recording conditions were carefully matched across species to ensure comparability, and a wide range of passive and active membrane properties was quantified to enable robust cross-species comparisons and aid interpretation of observed differences. To facilitate comparison with previously published studies, membrane potentials are reported both with and without correction for the liquid junction potential (LJP, 13 mV), because correction practices differ across laboratories and can complicate cross-study comparisons. We found that both resting membrane potential (RMP) and spontaneous firing rate (SFR) varied rhythmically across the LD cycle in mouse and fly neurons (Figure 2B, C and Table 1). RMP ranged from -47 to -71 mV (-34 to -58 mV LJP uncorrected) in VIP neurons and from -40 to -67 mV (-27 to -54 mV uncorrected) in l-LNvs. The amplitude of rhythmic variation was ±1.7 mV in l-LNvs and ±1.3 mV in VIP neurons, with a MESOR of -54 mV (-41 mV uncorrected) and -58 mV (-45 mV uncorrected), respectively, making l-LNvs on average about 4 mV more depolarised. These were modest but consistent rhythmic changes. Most neurons were spontaneously active, though 24% of VIP and 32% of l-LNv cells were silent under resting conditions. SFRs reached up to 8.4 Hz (VIP) and 7.2 Hz (l-LNv) and showed daily modulation with amplitudes of ±0.7 Hz (VIP) and ±0.5 Hz (l-LNv) around a MESOR of 1.8 Hz and 0.9 Hz, respectively. Thus, despite being slightly more hyperpolarised, VIP neurons consistently fired faster than l-LNvs across the entire 24h cycle (∼1 Hz on average). Sinusoidal fits with a 24 h period revealed that both parameters oscillated in phase, peaking around midday (acrophase ZT: RMP - VIP 6.5 h, l-LNv 5.4 h; SFR - VIP 6.1 h, l-LNv 5.1 h). Furthermore, RMP and SFR were positively correlated within each species, with comparable slopes of 0.13 Hz/mV (VIP) and 0.16 Hz/mV (l-LNv; Figure 2D and Table 1), indicating close voltage-to-firing coupling across species. Together, these findings show that mouse and fly clock neurons share strikingly similar electrophysiological dynamics: both are more depolarised and fire most rapidly around the middle of the day (midday) and become hyperpolarised and less active around the middle of the night (midnight). This alignment of membrane and firing rhythms across phyla highlights deeply conserved cellular mechanisms underlying circadian timekeeping.

**Figure 2:**
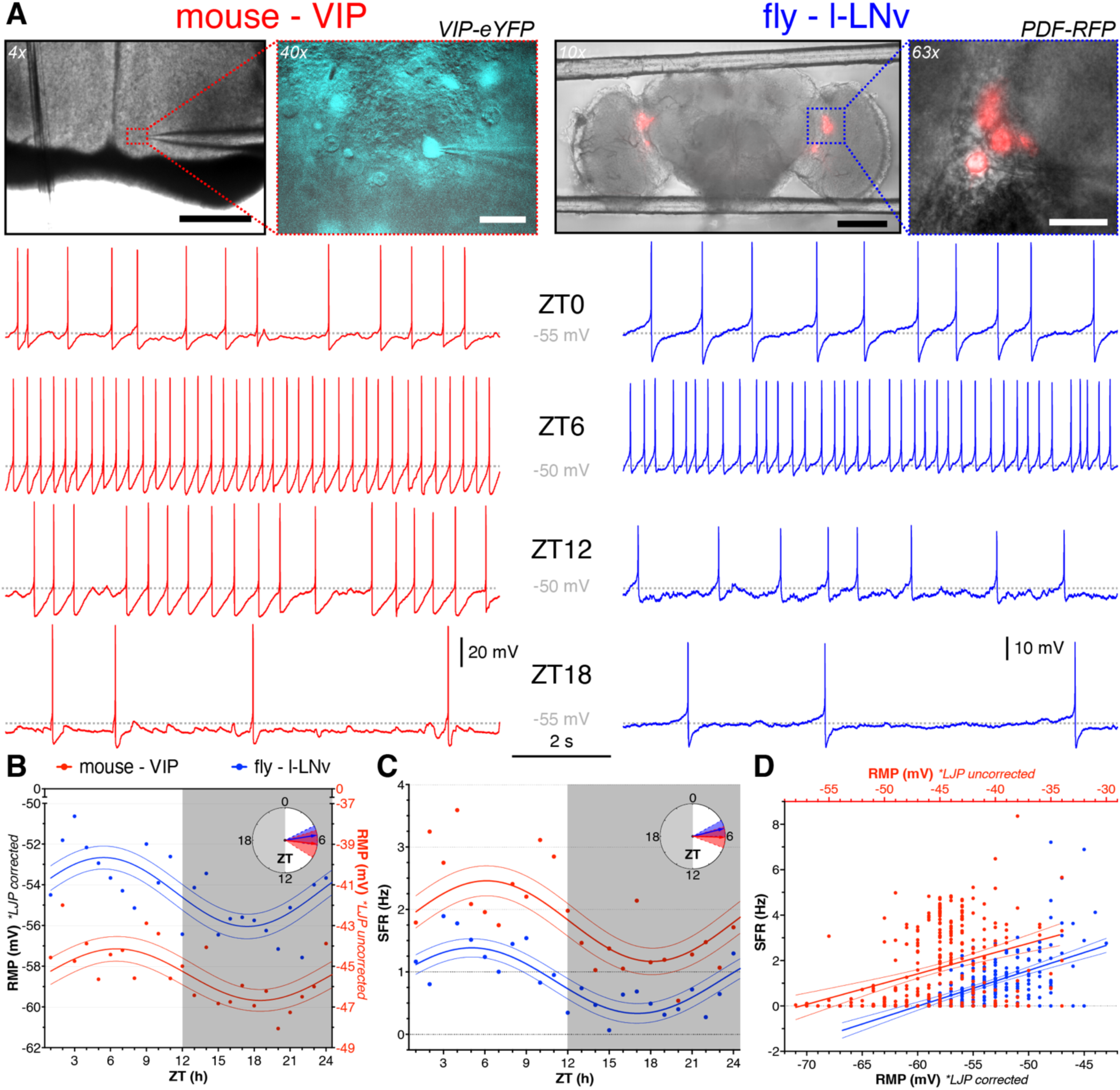
Diurnal variation in spontaneous electrical activity of mouse and fly clock neurons. (**A**) Bright-field and fluorescence images of whole-cell patch-clamp recordings from mouse (left) and fly (right) clock neurons. In mice, VIP-expressing neurons are located in the ventral SCN, just above the optic chiasm on either side of the third ventricle; scale bar, 500 µm. In flies, PDF-expressing large ventrolateral neurons (l-LNv) are situated on either side in a cleft separating the central brain from the medulla; scale bar, 100 µm. Inserts show patch pipettes targeting identified clock neurons; scale bars, 25 µm. Below, examples of current-clamp recordings from VIP (red) and l-LNv (blue) neurons at indicated Zeitgeber Times (ZTs), illustrating time-dependent differences in spontaneous electrical activity in firing neurons. (**B**, **C**) Diurnal profiles of resting membrane potential (RMP, **B**) and spontaneous firing rate (SFR, **C**) across 24 h under LD conditions. Symbols represent individual means for mouse (red) and fly (blue) recordings at each timepoint. Lines show the best fit and 95% confidence intervals of a 24-hour sine wave model. Clock face insets show phase distributions of peak RMP and SFR, with arrow direction indicating peak phase, length indicating fit strength, and shaded areas showing 95% confidence intervals. Both parameters peak near ZT6 in both species. n = 275 (VIP), n = 252 (l-LNv). (**D**) SFR and RMP are positively correlated in both species, indicating that more depolarised neurons tend to fire at higher rates. Lines show the best fit and 95% confidence intervals of a linear model. n = 302 (VIP), n = 261 (l-LNv). Both LJP corrected (black left and bottom axes) and uncorrected (red right and top axes) values are shown in (B) and (D). See Table 1 and Supplementary Figure 1 for details.

**Table 1:** Circadian curve-fitting and regression analysis of neurophysiological parameters in mouse VIP and fly l-LNv neurons.

| | mouse / fly | n | amplitude | acrophase<br>ZT [h] | MESOR | correct<br>probability | $\Delta AICc$ | R<sup>2</sup> |
| --- | --- | --- | --- | --- | --- | --- | --- | --- |
| RMP | VIP | 275 | 1.27 | 6.50 | -58.41 | >99.99% | 21.73 | 0.053 |
| (mV) | I-LNv | 252 | 1.69 | 5.42 | -54.36 | >99.99% | 36.00 | 0.091 |
| R<sub>in</sub> | VIP | 274 | 0.15 | 9.97 | 1.17 | >99.99% | 23.11 | 0.049 |
| (GΩ) | I-LNv | 244 | 0.38 | 18.96 | 2.59 | >99.99% | 35.03 | 0.091 |
| capacitance | VIP | 274 | 1.63 | 4.24 | 32.77 | 78.68% | 2.61 | 0.010 |
| (pF) | I-LNv | 244 | 3.14 | 5.18 | 33.69 | >99.99% | 18.78 | 0.045 |
| rheobase | VIP | 270 | 4.70 | 19.85 | 28.37 | 99.96% | 15.46 | 0.037 |
| (pA) | I-LNv | 220 | 2.70 | 7.67 | 25.33 | >99.99% | 20.31 | 0.058 |
| f at +30 pA | VIP | 272 | 0.70 | 4.54 | 10.13 | 91.75% | 4.82 | 0.014 |
| (Hz) | I-LNv | 237 | 0.55 | 4.10 | 22.19 | 46.60% | -0.27 | 0.004 |
| gain | VIP | 272 | 0.02 | 2.31 | 0.30 | 74.78% | 2.17 | 0.008 |
| (Hz/pA) | I-LNv | 238 | 0.03 | 3.77 | 0.62 | 89.50% | 4.29 | 0.017 |
| SFR | VIP | 275 | 0.65 | 6.12 | 1.81 | >99.99% | 34.66 | 0.084 |
| (Hz) | I-LNv | 252 | 0.53 | 5.05 | 0.86 | >99.99% | 55.43 | 0.124 |
| I<sub>h</sub> at −30 pA | VIP | 274 | 1.03 | 2.01 | 5.43 | 99.99% | 18.00 | 0.037 |
| (mV) | I-LNv | 239 | 1.76 | 20.74 | 5.09 | >99.99% | 21.64 | 0.061 |
| Linear regression analysis of SFR in relation to RMP |  |  |  |  |  |  |  |  |
|  | mouse / fly | n | slope | y-intercept | R<sup>2</sup> | slope non-zero | slopes different | elevations different |
| SFR | VIP | 302 | 0.134 | 9.47 | 0.127 | *p* < 0.001 | *p* = 0.37 | *p* < 0.001 |
| (Hz/mV) | I-LNv | 261 | 0.157 | 9.43 | 0.359 | *p* < 0.001 |  |  |
| Diurnal rhythms were assessed by comparing a 24-h sine-wave model with non-zero baseline to a linear model using Akaike's Information Criterion (ΔAICc); the correct probability indicates preference for the sine-wave model and goodness-of-fit is reported as R<sup>2</sup>. Amplitude is defined as the peak-to-MESOR difference; acrophase indicates the Zeitgeber time (ZT) of the fitted peak; MESOR denotes the midline estimating statistic of rhythm. RMP, resting membrane potential (+13 mV LJP-corrected baseline values shown); R<sub>in</sub>, input resistance; f at +30 pA, firing rate at +30 pA current injection; SFR, spontaneous firing rate; I<sub>h</sub> at -30 pA, h-current amplitude at -30 pA current injection. |  |  |  |  |  |  |  |  |
| Linear regression analyses relating SFR to RMP report goodness-of-fit (R<sup>2</sup>), tests for non-zero slopes, and between-species comparisons of slopes and elevations. |  |  |  |  |  |  |  |  |

### Divergent active and passive membrane properties across species

Having confirmed that both neuron types exhibit rhythmic membrane and firing properties, we next examined additional passive and active membrane properties to further characterise the electrophysiological profiles of l-LNv and VIP neurons and determine whether these features are similarly conserved or divergent across species. When depolarising current was injected, neurons depolarised and either initiated or increased their firing rate in proportion to the stimulus strength, with +30 pA serving as a standard reference current for comparison (Figure 3A). While l-LNvs typically fired repetitively throughout the entire depolarising pulse, a proportion of VIP neurons showed spike-frequency adaptation, slowing and ceasing firing before the end of the pulse (26%, 70 of 270 spiking cells), compared with only 1% (3 of 253) in l-LNvs. Overall, l-LNvs were more excitable than VIP neurons, firing at higher rates in response to the same +30 pA depolarising current (MESOR 22 Hz vs 10 Hz) and showed no diurnal variation, while VIP neurons showed only weak cycling (Figure 3C and Table 1).

**Figure 3:**
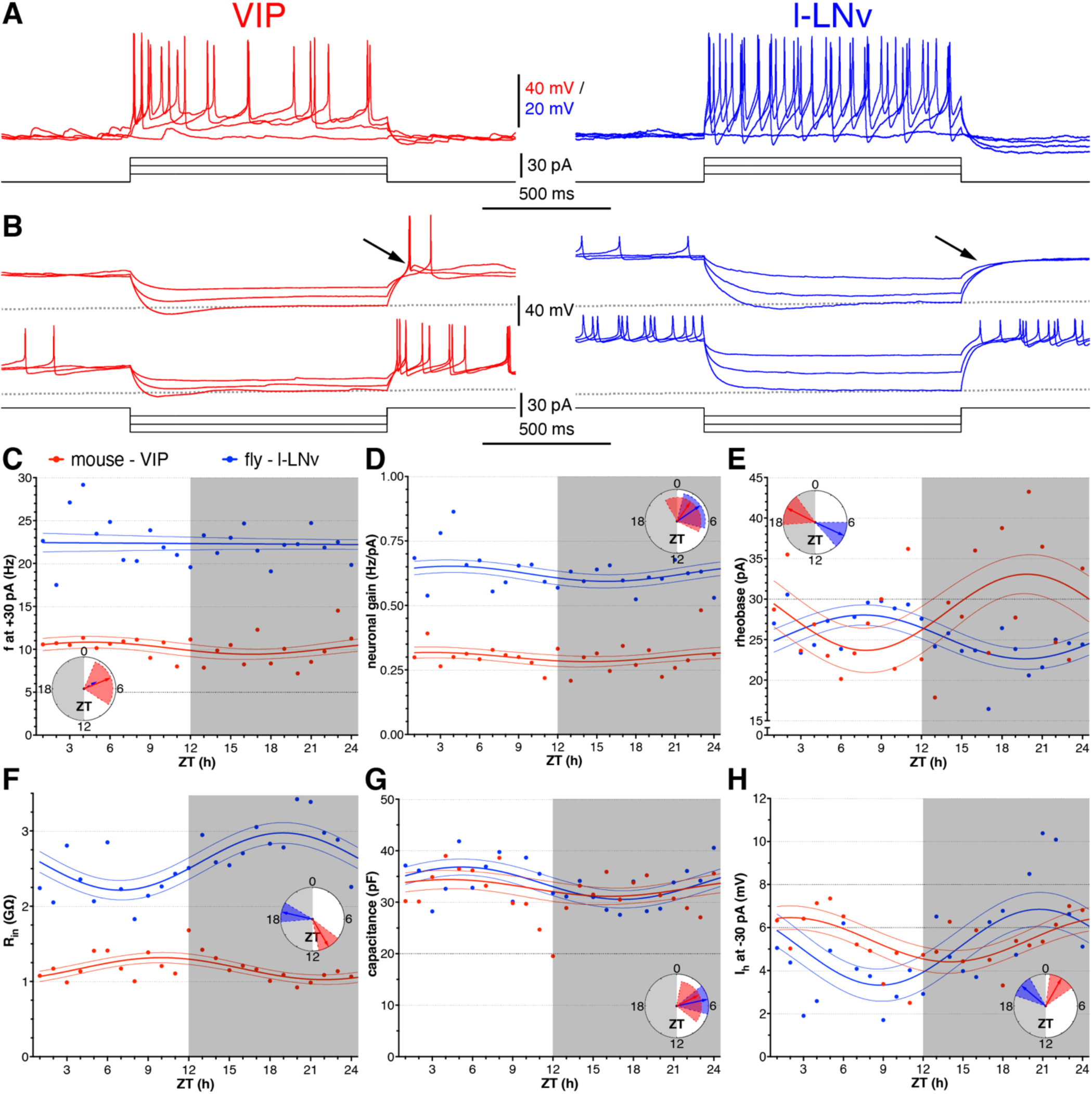
Diurnal variation in active and passive membrane properties in mouse and fly clock neurons. (**A**) Representative current-clamp recordings from VIP (red, left) and l-LNv (blue, right) neurons in response to depolarising current steps used to calculate firing rate at +30 pA and neuronal gain. (**B**) A couple of representative current-clamp recordings in response to hyperpolarising current steps used to calculate input resistance (R_in_) and h-currents (I_h_). Note the pronounced post-inhibitory rebound excitation (arrow) in VIP neurons (observed in 93% of recordings), which is absent in l-LNv (arrow). (**C**-**H**) Diurnal changes in firing rate at +30 pA (f at +30 pA, **C**), neuronal gain (**D**), rheobase (**E**), R_in_ (**F**), cell capacitance (**G**), and h-current (I_h_ at -30 pA, **H**) over 24 hours. Symbols show individual means (mouse: red; fly: blue). Lines represent 24 h sine fits with 95% confidence intervals. Clock insets indicate peak phases (arrow direction), fit strength (arrow length), and 95% confidence intervals (shaded arcs). Firing rate at +30 pA remains stable in fly l-LNv neurons but shows daily variation in VIP neurons. Neuronal gain peaks at a similar phase in both species, while rheobase shows opposite phases in mouse and fly. R_in_ rhythms are nearly in antiphase between species; capacitance peaks at a similar phase, while I_h_ shows opposing trends around ZT0. n = 272 (VIP), n = 237 (l-LNv) for C; n = 272 (VIP), n = 238 (l-LNv) for D; n = 270 (VIP), n = 220 (l-LNv) for E; n = 274 (VIP), n = 244 (l-LNv) for F, G; n = 274 (VIP), n = 239 (l-LNv) for H. See Table 1 and Supplementary Figure 1 for details.

The firing-current (f-I) relationship revealed a higher neuronal gain - the slope of the linear portion of the f-I curve, reflecting the sensitivity of firing rate to input strength - in l-LNvs (MESOR: VIP 0.3 Hz/pA, l-LNv 0.6 Hz/pA); with both cell types showing modest diurnal cycling that peaked in the morning (Figure 3D and Table 1). We next quantified the rheobase, the minimal current required to evoke an action potential, by holding neurons at -73 mV (-60 mV uncorrected) and injecting brief (50 ms) current steps. Rheobase values were similar between the two neuron types (MESOR: VIP 28 pA, l-LNv 25 pA) and showed daily variations but oscillated in antiphase, with l-LNvs peaking later in the day and VIP neurons later in the night (Figure 3E and Table 1).

Injection of hyperpolarising current caused the membrane potential to decrease to a stable plateau (Figure 3B). From the resulting current-voltage (I-V) relationship, we determined the input resistance (R_in_), which reflects passive membrane leak and is inversely related to membrane conductance; higher R_in_ indicates fewer open ion channels. R_in_ showed robust rhythmicity and was nearly in antiphase between cell types, with VIP neurons peaking in the day and exhibiting roughly half (MESOR: VIP 1.2 GΩ, l-LNv 2.6 GΩ) the R_in_ of l-LNvs that are peaking in the night (Figure 3F and Table 1). Cell capacitance, calculated from the membrane time constant (τ) and R_in_, provides a measure of the neuron’s effective membrane area. Capacitance values indicated that the two neuron types are of comparable size (MESOR: VIP 33 pF, l-LNv 34 pF) and that capacitance cycles in phase in both, peaking in the middle of the day (Figure 3G and Table 1). Some neurons displayed an initial sag during hyperpolarisation, indicative of the hyperpolarisation-activated current (I_h_; present in 91% of VIP neurons, 273 of 301, and 70% of l-LNvs, 176 of 250). This current also exhibited diurnal modulation, peaking around ZT0 but with slightly opposing phase relationships between species - before lights-on in l-LNvs and after lights-on in VIP neurons - while overall MESOR was comparable (VIP 5.4 mV, l-LNv 5.1 mV; Figure 3H and Table 1). Finally, upon release from hyperpolarisation, most VIP neurons depolarised beyond their resting membrane potential and fired action potentials even when previously silent (Figure 3B left, top trace), demonstrating a strong post-inhibitory rebound (PIR). This feature was observed in nearly all VIP neurons (93%, 279 of 301) but was completely absent in l-LNvs (0 of 255; Figure 3B), highlighting a major functional difference between the two species. The absence of PIR in l-LNvs despite comparable I_A_ amplitudes (see below) suggests additional ionic or morphological factors shape rebound excitability. Together, these analyses reveal both conserved and cell type-specific rhythmic modulations of intrinsic membrane properties that shape daily patterns of excitability across the two species.

### Ionic mechanisms underlying excitability rhythms

Since daily variation in potassium currents is implicated in circadian rhythms of clock neuron electrical activity, we next aimed to understand the ionic basis of these rhythmic and species-specific features. To do so, we performed voltage-clamp recordings from both neuron types at midday (ZT5-7) and midnight (ZT17-19), corresponding to the peak and trough phases of most neurophysiological parameters, to isolate and quantify the underlying potassium currents. First, we assessed the A-type potassium current (I_A_), a fast-activating, transient outward current that inactivates within tens of milliseconds and can strongly influence excitability and post-inhibitory rebound (PIR) responses. Recordings were performed in the presence of TTX to block sodium channels and sodium-dependent currents from masking I_A_. We isolated I_A_ by recording a series of depolarising voltage steps from a hyperpolarised holding potential and subtracting currents obtained using the same protocol after a depolarising pre-pulse (Figure 4A). This allowed measurement of both current amplitude and inactivation kinetics (τ_fast_ and τ_slow_) at the highest tested command potential of +60 mV (+73 mV LJP uncorrected, Figure 4B). To account for differences in neuron size, currents were normalised to cell capacitance to yield current density, enabling comparison of conductances on a per-membrane-area basis. These experiments revealed that VIP neurons exhibited a larger I_A_ at midnight compared to midday (midday 8 pA/pF, midnight 23 pA/pF), whereas l-LNvs showed no significant diurnal difference (midday 15 pA/pF, midnight 18 pA/pF). Although I_A_ was larger at midnight in VIP neurons, overall amplitudes and decay kinetics (τ_fast_ and τ_slow_) were broadly comparable between cell types and time points (Figure 4C-E and Table 2).

**Figure 4:**
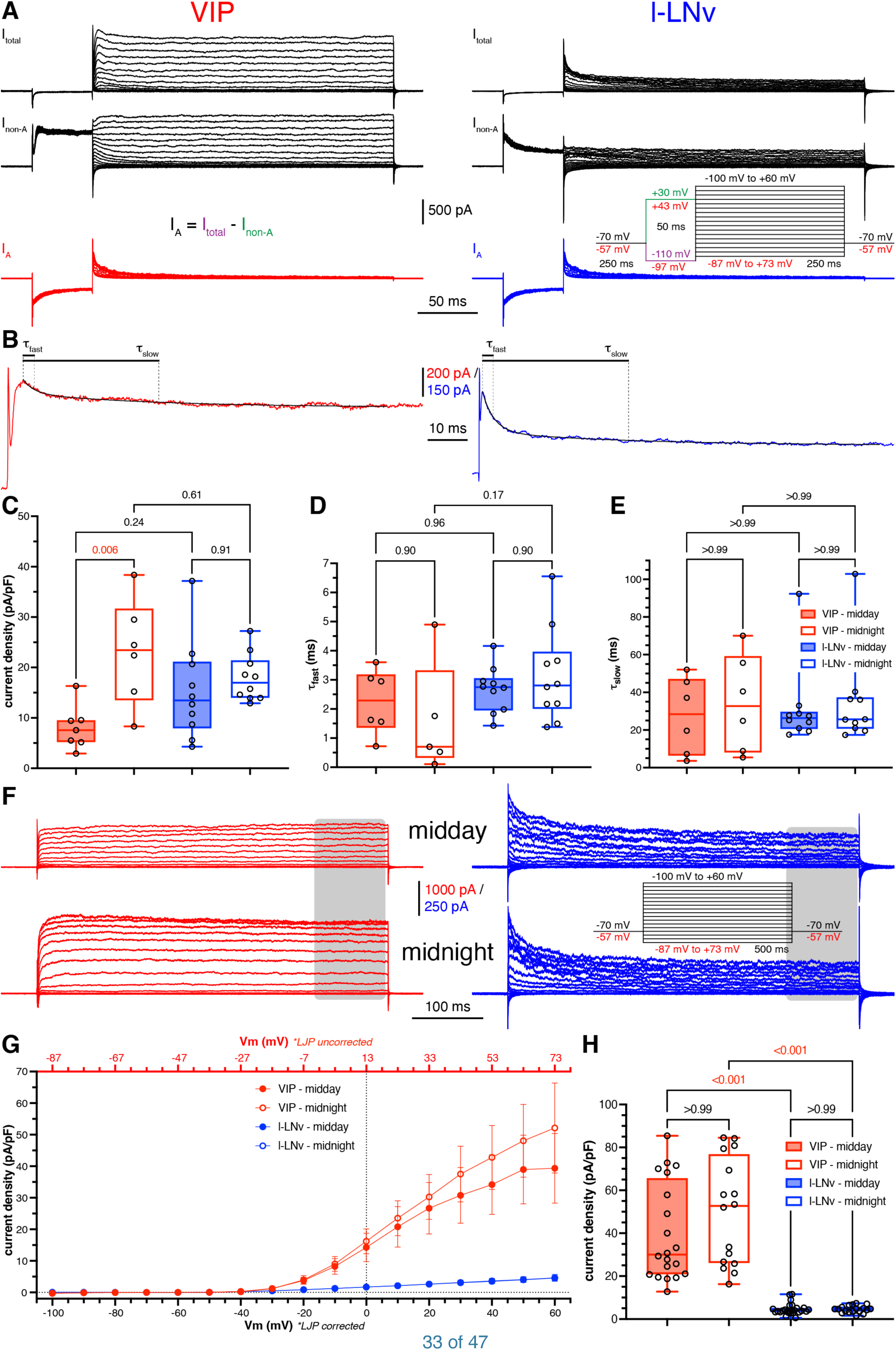
Voltage-clamp recordings of A-type and sustained potassium currents in mouse and fly clock neurons at midday and midnight. (**A**) Representative voltage-clamp traces from VIP (left) and l-LNv (right) neurons recorded in the presence of TTX. Top: Depolarising voltage steps from a holding potential of -110 mV (-97 mV LJP uncorrected) evoke the full potassium current (I_total_). Middle: The same protocol from a holding potential of +30 mV (+43 mV uncorrected) isolates potassium currents lacking the A-type component (I_non-A_). Bottom (coloured): Digital subtraction of I_non-A_ from I_total_ reveals the A-type current (I_A_). Stimulation protocol shown in inset. (**B**) Fast and slow inactivation time constants (τ_fast_ and τ_slow_) were derived from a double-exponential fit (black line) applied to I_A_ evoked by a depolarising step to +60 mV (+73 mV uncorrected) (expanded timescale). (**C**-**E**) Quantification of I_A_ current density at +60 mV (+73 mV uncorrected) (**C**), τ_fast_ (**D**) and τ_slow_ (E) shows slightly larger VIP A-type currents at midnight compared to midday but otherwise not much diurnal variation. (**F**) Representative voltage-clamp traces from VIP (red, left) and l-LNv (blue, right) neurons recorded at ZT6 (midday, top) and ZT18 (midnight, bottom) in response to a series of hyperpolarising to depolarising voltage steps (protocol shown in inset). Sustained currents were measured at the end of each voltage step (indicated by grey bars). (**G**) Current-voltage (I-V) relationships reveal higher current densities in VIP neurons compared to l-LNv neurons, with both showing a similar onset of the sustained current around -40 mV (-27 mV uncorrected). Circles, means; error bars, 95% CI. (**H**) Quantification of current density at +60 mV (+73 mV uncorrected) shows no significant time-of-day differences in either species. Bars, means; error bars, 95% CI; *p*-values indicated. See Table 2 for details.

**Table 2:** Statistical details for voltage-clamp experiments and sinusoidal stimulations.

|  | mouse / fly | mean | 95% CI | n | midday vs<br>midnight | mouse vs fly | stat. test |
| --- | --- | --- | --- | --- | --- | --- | --- |
| voltage-clamp experiments |  |  |  |  |  |  |  |
| I<sub>A</sub> (pA/pF) | VIP | 8.07 | 4.03 | 7 |  | p = 0.24 | ANOVA<br>with Sidak's |
| midday | I-LNv | 15.33 | 6.96 | 10 |  |  |  |
| I<sub>A</sub> (pA/pF) | VIP | 23.04 | 11.08 | 6 | p = 0.006 | p = 0.61 | KW with<br>Dunn's |
| midnight | I-LNv | 17.94 | 3.36 | 10 | p = 0.91 |  |  |
| τ<sub>fast</sub> (ms) | VIP | 2.25 | 1.17 | 6 |  | p = 0.96 | ANOVA<br>with Sidak's |
| midday | I-LNv | 2.67 | 0.56 | 10 |  |  |  |
| τ<sub>fast</sub> (ms) | VIP | 1.60 | 2.41 | 5 | p = 0.90 | p = 0.17 | ANOVA<br>with Sidak's |
| midnight | I-LNv | 3.15 | 1.15 | 10 | p = 0.90 |  |  |
| τ<sub>slow</sub> (ms) | VIP | 27.51 | 21.32 | 6 |  | p > 0.99 | KW with<br>Dunn's |
| midday | I-LNv | 31.64 | 15.62 | 10 |  |  |  |
| τ<sub>slow</sub> (ms) | VIP | 34.17 | 27.15 | 6 | p > 0.99 | p > 0.99 | KW with<br>Dunn's |
| midnight | I-LNv | 34.86 | 17.99 | 10 | p > 0.99 |  |  |
| I<sub>sus</sub> (pA/pF) | VIP | 39.94 | 10.66 | 20 |  | p < 0.001 | KW with<br>Dunn's |
| midday | I-LNv | 4.56 | 1.05 | 26 |  |  |  |
| I<sub>sus</sub> (pA/pF) | VIP | 49.88 | 13.14 | 16 | p > 0.99 | p < 0.001 | KW with<br>Dunn's |
| midnight | I-LNv | 4.56 | 8.81 | 18 | p > 0.99 |  |  |
| sinusoidal stimulations |  |  |  |  |  |  |  |
| spikes at 1 Hz | VIP | 10 |  | 223 |  |  |  |
|  | I-LNv | 10 |  | 201 |  |  |  |
| spikes at 2 Hz | VIP | 5.55 | 0.16 | 223 |  | p < 0.001 | 2-way<br>ANOVA |
|  | I-LNv | 4.81 | 0.08 | 201 |  |  |  |
| spikes at 5 Hz | VIP | 2.90 | 0.14 | 223 |  | p < 0.001 | 2-way<br>ANOVA |
|  | I-LNv | 2.12 | 0.07 | 201 |  |  |  |
| spikes at 10 Hz | VIP | 1.47 | 0.10 | 223 |  | p < 0.001 | 2-way<br>ANOVA |
|  | I-LNv | 1.05 | 0.05 | 201 |  |  |  |
| KW, Kruskal-Wallis with Dunn's *post hoc* test. |  |  |  |  |  |  |  |

We then measured sustained (delayed rectifier) potassium currents, which activate more slowly and do not inactivate, as assessed at the end of a long depolarising voltage step (Figure 4F). These currents contribute to membrane repolarisation after action potentials, regulate firing frequency, and stabilise the resting potential, thus shaping overall neuronal excitability. Sustained currents began to activate around -40 mV (-27 mV uncorrected) in both cell types and increased nearly linearly up to the highest tested voltage of +60 mV (+73 mV uncorrected, Figure 4G). Despite similar voltage dependence, their amplitude differed markedly between species, being approximately ten times larger in VIP neurons (midday: VIP 40 pA/pF, l-LNv 5 pA/pF; midnight: VIP 50 pA/pF, l-LNv 5 pA/pF). This difference could only partly be explained by the approximately twofold higher input resistance in l-LNv, which would be expected to reduce macroscopic potassium currents by a similar factor, suggesting that other factors, such as reduced channel density or space clamp limitations, may also contribute. No significant diurnal modulation of these sustained currents was observed in either cell type (Figure 4H and Table 2). Together, these voltage-clamp recordings indicate that while sustained potassium currents differ greatly between species, diurnal modulation of both A-type and sustained currents is only modest.

### Frequency-dependent responsiveness and temporal fidelity

Having established the basic ionic currents, we next tested how these properties translate into information processing by probing frequency-dependent responses and temporal fidelity, i.e. how neurons respond to inputs of varying frequency. Cells were hyperpolarised to - 73 mV (-60 mV LJP uncorrected), and sinusoidal current waveforms of increasing frequency (10 cycles each at 1, 2, 5, and 10 Hz) were injected, with current amplitudes adjusted to elicit approximately five spikes at the highest frequency (10 Hz). Both VIP and l-LNv neurons fired more action potentials at slower stimulation rates, indicating reduced temporal summation at higher frequencies (Figure 5A). Heat maps of normalised firing responses (relative to maximal spiking) revealed no diurnal modulation in either species, with similar responses across the 24 h cycle (Figure 5B). Notably, response patterns differed between species during the first stimulus cycle: VIP neurons exhibited strong initial firing followed by a steady plateau across all tested frequencies, whereas l-LNvs showed lower initial firing before reaching a similar plateau (Figure 5C). Both neuron types displayed a roughly twofold reduction in spiking as stimulation frequency increased from 1 to 2 to 5 to 10 Hz, with VIP neurons maintaining slightly higher firing rates overall (Figure 5D and Table 2). These results indicate that both mouse and fly neurons can reliably follow and sustain firing across all tested frequencies, showing no clear frequency preference or filtering within the 1-10 Hz range tested, and thus responding broadly to input rates within this range.

**Figure 5:**
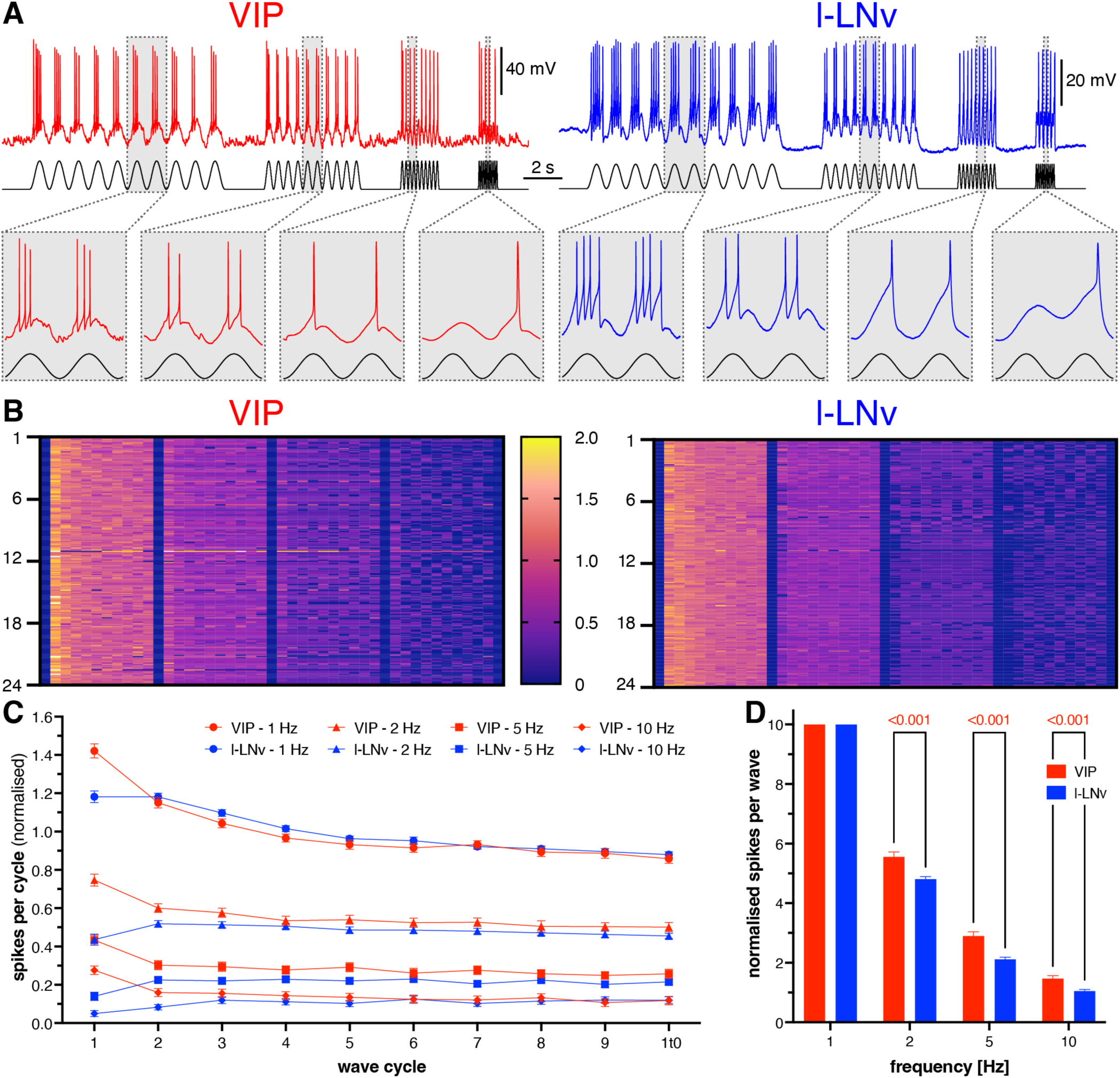
Frequency-dependent firing of mouse and fly clock neurons in response to sinusoidal stimulation. (**A**) Representative current-clamp recordings from VIP (red, left) and l-LNv (blue, right) neurons during sinusoidal current stimulation. Each train consisted of 10 cycles delivered at increasing frequencies (1 Hz, 2 Hz, 5 Hz, 10 Hz). Insets (below) show two expanded cycles per frequency at a slower timescale. (**B**) Heatmap of normalised spike responses per cycle across the four stimulation frequencies, scaled to 10 spikes at 1 Hz (i.e. one per cycle), organised by ZT. Responses were independent of ZT, with higher firing at lower stimulation frequencies, but no evidence of resonance. (**C**) Normalised spike number per individual cycle across stimulation frequencies, showing a strong response to the first cycle, followed by a gradual decline for VIP neurons. In contrast, l-LNv neurons exhibit lower initial firing before the plateau. Symbols, means; error bars, 95% CI. (**D**) Average normalised number of spikes per 10-cycle train at each frequency. Spike counts decrease with increasing frequency in both species, with l-LNv neurons showing relatively fewer spikes than VIP neurons. Bars, means; error bars, 95% CI. n = 223 (VIP), n = 201 (l-LNv). See Table 2 for details.

In summary, circadian neurons from mouse and fly share conserved daily rhythms in key electrophysiological properties despite species-specific differences reflecting their temporal niches. Resting membrane potential, firing rate, and neuronal gain oscillated with similar phases, whereas input resistance, rheobase, and post-inhibitory rebound showed divergent timing or presence. Although ionic currents differed in magnitude - particularly sustained potassium currents in VIP neurons - which would normally act to reduce excitability, both neuron types nevertheless maintained comparable firing responses and reliably followed inputs across frequencies. These results show that evolutionarily distant clock neurons use partially divergent mechanisms to achieve conserved temporal coding, highlighting both conservation and flexibility in the cellular basis of circadian timekeeping.

## Discussion

In this study, we investigated the membrane hand of the circadian clock in mice and flies to understand how conserved and divergent electrophysiological mechanisms shape rhythmic neuronal function in evolutionarily distant species occupying different temporal niches. Although mouse SCN VIP and *Drosophila* l-LNv PDF neurons occupy analogous positions within their respective circadian circuits - serving key roles in entrainment, timekeeping, and time-of-day signalling - the circuits they inhabit differ dramatically in scale, architecture, and environmental demands (Antle and Silver, 2005; Parisky *et al*., 2008; Jones *et al*., 2018; Helfrich-Förster and Reinhard, 2025). The *Drosophila* clock comprises ∼240 neurons and integrates both cryptochrome-based and visual photoreception, whereas the mammalian SCN contains ∼20,000 neurons and relies exclusively on retinal input for light entrainment. Likewise, behavioural adaptations such as diurnality in flies and nocturnality in mice impose distinct pressures on how individual clock neurons encode and distribute time-of-day information (Figure 1). Although light can exert both excitatory and inhibitory effects depending on pathway and state, daytime illumination constitutes a dominant modulatory drive that strongly shapes clock neuron excitability in diurnal flies (Sheeba *et al*., 2008b; Buhl *et al*., 2016; Schlichting *et al*., 2016; Tabuchi *et al*., 2021). In both species, VIP and PDF neurons function not only as intrinsic oscillators but as paracrine signalling hubs that broadcast timing signals, synchronise brain-wide clock networks and integrate photic and behavioural state information (Hegazi *et al*., 2019). Against this backdrop, our cross-species comparison reveals a mixture of conserved membrane features, reflecting shared computational requirements of circadian pacemakers, alongside clear species- and niche-specific adaptations in excitability and ionic mechanisms. Thus, despite occupying comparable network positions, VIP and l-LNv neurons exhibit electrophysiological specialisations that may reflect the differing circuit architectures and ecological constraints under which each clock operates.

We found that the most fundamental membrane properties of clock neurons are strikingly conserved across evolution and temporal niche. Both VIP and l-LNv neurons showed robust daily rhythms in resting membrane potential (RMP) and spontaneous firing rate (SFR) that were remarkably in phase (peak at midday) while modest in amplitude (∼3 mV and ∼1 Hz; Figure 2). Given the inherent challenges of cross-species comparisons, we carefully calibrated our experiments to ensure that measurements were as directly comparable and reliable as possible. This included maintaining roughly equal sample sizes across all Zeitgeber-time points to avoid phase bias, strictly avoiding light exposure during dark-phase dissections, and closely matching recording conditions across species, with solutions, protocols, equipment, and procedures kept as similar as possible between mouse and fly preparations. As a result, we were able for the first time to obtain precise and directly comparable measurements of RMP and SFR rhythms in circadian neurons from exemplar vertebrate and invertebrate species. These values closely match prior reports in both flies (Park and Griffith, 2006; Sheeba *et al*., 2008a; Buhl *et al*., 2016; Curran *et al*., 2019) and mice (Hermanstyne *et al*., 2016; Patton *et al*., 2020; Yang *et al*., 2023). Larger amplitudes reported in some studies may reflect differences in recording conditions, sampling across heterogeneous clock neuron populations, or limited cell-type identification - particularly in earlier work where recordings were not restricted to defined subsets such as VIP neurons - as well as variation in zeitgeber timing (Kuhlman and McMahon, 2004; Belle *et al*., 2009; Flourakis *et al*., 2015). Although the l-LNv peak phase we observe is somewhat later than previously described (Cao and Nitabach, 2008), it aligns well with population-level activity markers: Ca²⁺ imaging shows peak l-LNv activity around midday (Liang *et al*., 2016; Liang *et al*., 2017).

Beyond intrinsic electrical rhythms, circadian neurons also exhibit pronounced daily structural remodelling that can directly shape membrane properties. Both small LNvs in flies (Fernández *et al*., 2008) and VIP neurons in mice (Neitz *et al*., 2024) show diurnal changes in neurite complexity, with more extensive processes during the day, and l-LNv neurons likely follow a similar pattern, as their synaptic terminals decrease during sleep and increase after sleep deprivation (Donlea *et al*., 2009; Inami and Koh, 2024). These morphological changes have functional consequences. An electron microscopy study revealed daily plasticity of presynaptic varicosities modulates the influence of clock neurons within the network (Ispizua *et al*., 2025). Moreover, reciprocal anti-phase remodelling within the s-LNvs and DN1s inhibitory network reconfigures network topology and modulates behavioural outputs such as light-induced startle responses, which are stronger at night when light is unexpected (Song *et al*., 2021). Consistent with this structural plasticity, we observed daily rhythms in cell capacitance in both neuron types, with similar phase and amplitude (Figure 3). Despite their markedly different anatomical arborisations, somatically measured capacitance was comparable, indicating that these rhythms reflect relative structural remodelling over the circadian cycle rather than differences in absolute morphological scale. Notably, l-LNvs possess far more extensive arborisations, including bilateral projections into the medulla. Such complex morphology is likely to impose space-clamp limitations during somatic voltage-clamp recordings, leading to an underestimation of ionic currents originating in distal processes. Despite this potential limitation, l-LNvs nonetheless remain more excitable than VIP neurons, reinforcing the conclusion that their heightened excitability reflects genuine physiological differences rather than recording artefacts. Daily changes in capacitance are therefore likely to reflect underlying morphological remodelling, which would influence input resistance (R_in_) and membrane time constant (τ), and thus reshape integration windows over the circadian cycle.

Input resistance varied across the day and showed pronounced species-specific differences: in line with previous reports (Park and Griffith, 2006; Cao and Nitabach, 2008; Hermanstyne *et al*., 2016; Buhl *et al*., 2019; Yang *et al*., 2023), l-LNvs had roughly double the R_in_ of VIP neurons, and our data reveal that these rhythms are temporally inverted, peaking in antiphase (Figure 3). As capacitance was similar between cell types, the higher R_in_ in l-LNvs produced a correspondingly longer τ, as confirmed by our measurements. This combination predicts slower membrane charging and larger voltage deflections in response to a given synaptic or injected current. At the level of ionic mechanisms, the higher R_in_ of l-LNvs was accompanied by substantially smaller sustained outward currents (I_sus_) in voltage clamp, indicating lower overall membrane conductance. Strikingly, the amplitude and kinetics of the A-type potassium current (I_A_), a transient voltage-gated K^+^ current that rapidly activates and inactivates to delay action potential firing and shape membrane responsiveness, were comparable between species (Figure 4). As a result, I_A_ exerts a relatively greater influence on the overall membrane behaviour of l-LNvs, where passive conductance is lower. The reduced leak conductance in l-LNvs is also likely to contribute to their slightly more depolarised resting membrane potential. Functionally, these properties position l-LNvs as high-gain elements within a sparse circuit: in a network comprising only a handful of PDF neurons, elevated R_in_ would enhance the salience of weak or sparse synaptic and photic inputs. The daytime reduction in R_in_ may therefore act as a form of gain control, limiting over-responsiveness to sustained light drive while preserving sensitivity to behaviourally relevant signals. Together, these findings indicate that species differences in excitability arise primarily from the balance of passive membrane properties and outward conductances, rather than from changes in the basic kinetics of fast voltage-gated channels.

Interestingly, rheobase also cycled in anti-phase, reaching a trough during the respective behavioural inactive phase in each species - daytime for mice and midnight for flies. In l-LNvs, these R_in_ and rheobase rhythms were dissociated from RMP cycles: typically, higher R_in_ coincides with a more depolarised baseline and lower rheobase (Carrascal *et al*., 2011), yet here the phases did not consistently align. This suggests that mechanisms controlling baseline membrane potential and dynamic excitability, the neuron’s responsiveness to depolarising currents, are at least partially decoupled, allowing distinct tuning of neuron responsiveness across the circadian cycle. Differences in spike output further highlight divergent excitability strategies. Despite slightly more hyperpolarised RMPs, VIP neurons fired at ∼1 Hz higher baseline rates; however, this was not due to lower action potential threshold or intrinsic excitability, as rheobase was similar and l-LNvs produced more spikes in response to depolarisation. Lower excitability in VIP neurons - fewer spikes per current and some silent cells - likely reflects stronger regulation by voltage-gated conductances such as SK, BK, or sodium-channel inactivation due to depolarisation blockade (Belle *et al*., 2009; Pierre-Ferrer *et al*., 2024). These patterns suggest that clock neurons tune excitability according to network constraints: VIP neurons, embedded in a large and redundant population (∼2200 cells), prioritise stability and controlled responsiveness, whereas the sparse population of l-LNvs (only 8 cells) requires higher intrinsic gain and slower integration to ensure reliable signal propagation within the circuit.

A striking species difference was the presence of prominent post-inhibitory rebound (PIR) firing in VIP neurons and its complete absence in l-LNvs (Figure 3). PIR is a well-established mechanism for neuronal synchronisation and rhythm generation (e.g., in thalamocortical circuits; (Krosigk von *et al*., 1993) or central pattern generation (Marder and Bucher, 2001)) and is widespread in the SCN (Colwell, 2011), where it is thought to support rapid phase alignment and coupling among clock neurons. Its absence in l-LNvs therefore points to fundamentally different mechanisms of neuronal synchronisation and information integration in fly circadian circuits. Importantly, PIR is not generally absent from *Drosophila*, having been demonstrated in antennal lobe interneurons (Nagel and Wilson, 2016) and descending motor neurons (Roemschied *et al*., 2023), but it has not previously been reported in any fly clock neuron group. Several mechanisms could underlie the PIR in VIP neurons. Hyperpolarisation-activated cyclic nucleotide-gated (HCN) channels conducting I_h_ can depolarise the membrane following inhibition and prime neurons for rebound spiking; consistent with this, we detected I_h_ in most VIP neurons, in line with previous reports in the SCN (Atkinson *^e^t al.*, 201^1^; Colwell, 2011). PIR may also involve T-type Ca^2+^ channels, which de-inactivate during hyperpolarisation and generate transient inward currents upon release from inhibition, and are expressed in the SCN (McNally *et al*., 2020). In addition, network mechanisms are likely to contribute: SCN neurons are interconnected via reciprocal inhibition (Welsh *et al*., 2010), and GABAergic signalling in VIP neurons can be depolarising in a time-of-day-dependent manner due to circadian regulation of intracellular chloride (Klett *et al*., 2022). Together, these intrinsic and circuit-level features suggest that PIR in VIP neurons emerges from their interaction, positioning rebound firing as a powerful synchronising mechanism within the densely interconnected SCN. In contrast, l-LNvs express all these components associated with rebound firing yet nevertheless lack PIR. I_h_ is present in most l-LNvs at amplitudes comparable to VIP neurons and cycles across the day, peaking at night alongside maximal input resistance - a configuration that would ordinarily favour rebound excitability. However, in *Drosophila* clock neurons I_h_ might be functionally specialised: in s-LNvs it is required for high-frequency bursting and timed neuropeptide release (Fernandez-Chiappe *et al*., 2021), suggesting that I_h_ may preferentially support output signalling rather than synchronising rebound firing in the fly clock. *Drosophila* clock neurons also express the T-type Ca^2+^ channel homologue Ca-α1T, which is required for circadian output (Jeong *et al*., 2015; Liang *et al*., ^2^02^2^), and l-LNvs express GABA_A_ receptors (Parisky *et al*., 2008; Chung *et al*., 2009) within a reciprocally inhibitory clock network (Helfrich-Förster and Reinhard, 2025) that exhibits daily shifts in E_GABA_ (Buhl *et al*., 2016; Eick *et al*., 2022; Schellinger *et al*., 2022). The absence of PIR therefore cannot be explained by a simple lack of rebound-generating machinery but instead points to divergent circuit-level deployment of shared ionic components. A key difference appears to lie in the relative weighting of outward currents. In l-LNvs, the A-type potassium current (I_A_) is much larger relative to sustained outward currents compared to VIP neurons (Figure 4), a configuration that could suppress rebound excitation. Consistent with this, reducing I_A_ increases firing and excitability in both SCN neurons and l-LNvs (Hermanstyne *et ^a^l.*, 201^7^; Feng *et al*., 2018; Smith *et al*., 2019), and experimental and modelling work in the diurnal rodent *Rhabdomys pumilio* directly implicate I_A_ in shaping or delaying PIR (Bano-Otalora *et al*., 2021). Notably, I_A_ amplitude in l-LNvs was stable across the day, suggesting a constant brake on rebound firing, whereas VIP neurons showed time-of-day variation in I_A_. Together, this suggests that PIR is actively constrained in fly clock neurons, likely to prevent excessive excitation under continuous or prolonged photic drive, whereas mouse SCN neurons retain PIR to amplify and synchronise responses to sparse nocturnal cues. Rather than relying on rebound-mediated synchronisation, PDF neurons may achieve network coherence through sustained excitability, gain modulation, and paracrine peptidergic signalling within a smaller, more tightly coupled circuit. This divergence highlights how ecological context and network architecture shape the functional deployment of shared ion-channel repertoires in circadian systems.

Frequency-dependent firing revealed additional specialisations (Figure 5). Both VIP and l-LNv neurons exhibited broadband responsiveness, encoding low-frequency inputs faithfully while attenuating high-frequency signals. Spike output decreased roughly twofold when input frequency increased from 1 to 10 Hz, reflecting intrinsic membrane filtering rather than selective tuning to specific temporal features. VIP neurons displayed a slightly more adapting initial response, which could make them more sensitive to transient inputs, whereas the slower response of l-LNvs may arise from their prominent A-type potassium current. These frequency-dependent input-output relationships were time-of-day-independent, indicating that clock-state modulation primarily adjusts baseline excitability without compromising temporal fidelity of incoming signals. The ability to track inputs up to ∼10 Hz likely supports synchronisation among circadian neurons and reliable propagation of slow modulatory signals (VIP, PDF) without distortion. Notably, these neurons do not act as selective filters for temporal features, unlike many sensory or cortical neurons (Abbott and Regehr, 2004; Cariani and Baker, 2025); instead, they function as integrators of sustained drive, consistent with their role in long-timescale behavioural regulation. In this context, the larger but slower voltage responses of l-LNvs confer higher gain - reducing the amount of synaptic input required to elicit firing - while simultaneously acting as a low-pass filter limiting sensitivity to rapid fluctuations, a general principle of input-output transformations due to intrinsic conductances and passive membrane properties (Reyes, 2001). This trade-off favours temporal summation over fast tracking, stabilising firing output under persistent or noisy sensory conditions such as continuous daylight, at the cost of reduced responsiveness to brief, high-frequency inputs.

Importantly, not all measured parameters were rhythmic: evoked firing responses, sustained potassium currents, and frequency-dependent responses remained largely stable across the day/night cycle. This indicates that circadian modulation acts within a stable conductance framework that preserves reliable signal processing, while allowing modest, time-of-day-dependent fine-tuning of excitability. Apparent conductance rhythms likely arise from coordinated changes in ion-channel regulation and structural plasticity, with the summation of multiple coordinated small-amplitude fluctuations sufficient to shape neuronal responsiveness. Such slow, graded modulation reflects a design principle in which baseline excitability is actively stabilised, enabling precise temporal encoding while rhythmic shifts in RMP and ionic conductances adjust responsiveness rather than switching neurons on or off, aligning with the circadian clock’s role as an hours-scale integrator. Accordingly, circadian neurons function as low-pass, non-selective integrators specialised for slow-timescale homeostatic signalling, prioritising stable network communication over precise spike timing. Despite relying on distinct combinations of passive and active membrane properties, VIP and l-LNv neurons converge on remarkably similar daily electrical profiles, exemplifying functional convergence despite mechanistic divergence and illustrating how circadian systems separate slow excitability rhythms from robust, time-invariant encoding of synaptic input.

An important next step will be to determine which elements of the circadian membrane hand are cell-intrinsic and which depend on network interactions. This will require targeted perturbations combined with computational modelling. Notably, preliminary observations suggest species differences in the TTX sensitivity of resting membrane potential rhythms, raising the possibility that intrinsic mechanisms contribute more strongly to baseline excitability in l-LNvs than in VIP neurons. To establish how general these principles are, it will be essential to extend this framework to other *Drosophila* (s-LNvs, LNds, DN1s) and mouse SCN (AVP- or somatostatin/neurotensin-expressing) clock neuron classes. A further species-specific question concerns the interaction between light signalling and membrane excitability. In flies, cryptochrome (Cry) is directly light sensitive - a feature absent in mammals - and future studies should address how cryptochrome-dependent phototransduction interfaces with PDF neuron excitability during photic stimulation and entrainment (Fogle *et al*., 2011). Finally, circadian modulation of excitability may not be mediated exclusively by ion channels. Ion transporters such as NKCC are rhythmically regulated in both fly and mammalian clock neurons and play a prominent role in shaping intracellular chloride and GABAergic signalling (Alamilla *et al*., 2014; Buhl *et al*., 2016; Albers *et al*., 2017; Eick *et al*., 2022). Although these transporters have classically been considered electrically neutral, recent evidence that insect NKCC can generate a depolarising Na⁺ influx (Yarcusko *et al*., 2024) raises the possibility that electrogenic transport processes contribute directly to circadian membrane rhythms. Exploring such non-canonical mechanisms may therefore reveal an additional layer through which molecular clocks tune neuronal excitability.

In conclusion, the membrane hand of the circadian clock acts as the computational interface, the biophysical layer that translates intracellular molecular rhythms into changes in excitability level, enabling conserved functional outputs to emerge from divergent cellular mechanisms. In this framework, VIP neurons, through high conductance, low gain, adaptation and post-inhibitory rebound, may stabilise rhythmic synchrony and prevent runaway excitation, whereas l-LNv PDF neurons, with lower conductance but sustained excitability and higher gain, may support sustained output and controlled responsiveness to prolonged light input. Together, these cell-type-specific strategies exemplify computational degeneracy: distinct combinations of passive and active membrane properties achieve similar circadian functions despite substantial differences in circuit architecture and ecological context. This flexibility allows evolution to preserve core computational goals while tuning the underlying ionic machinery to meet species-specific behavioural and environmental demands.

## Figures

**Supplementary Figure 1:**
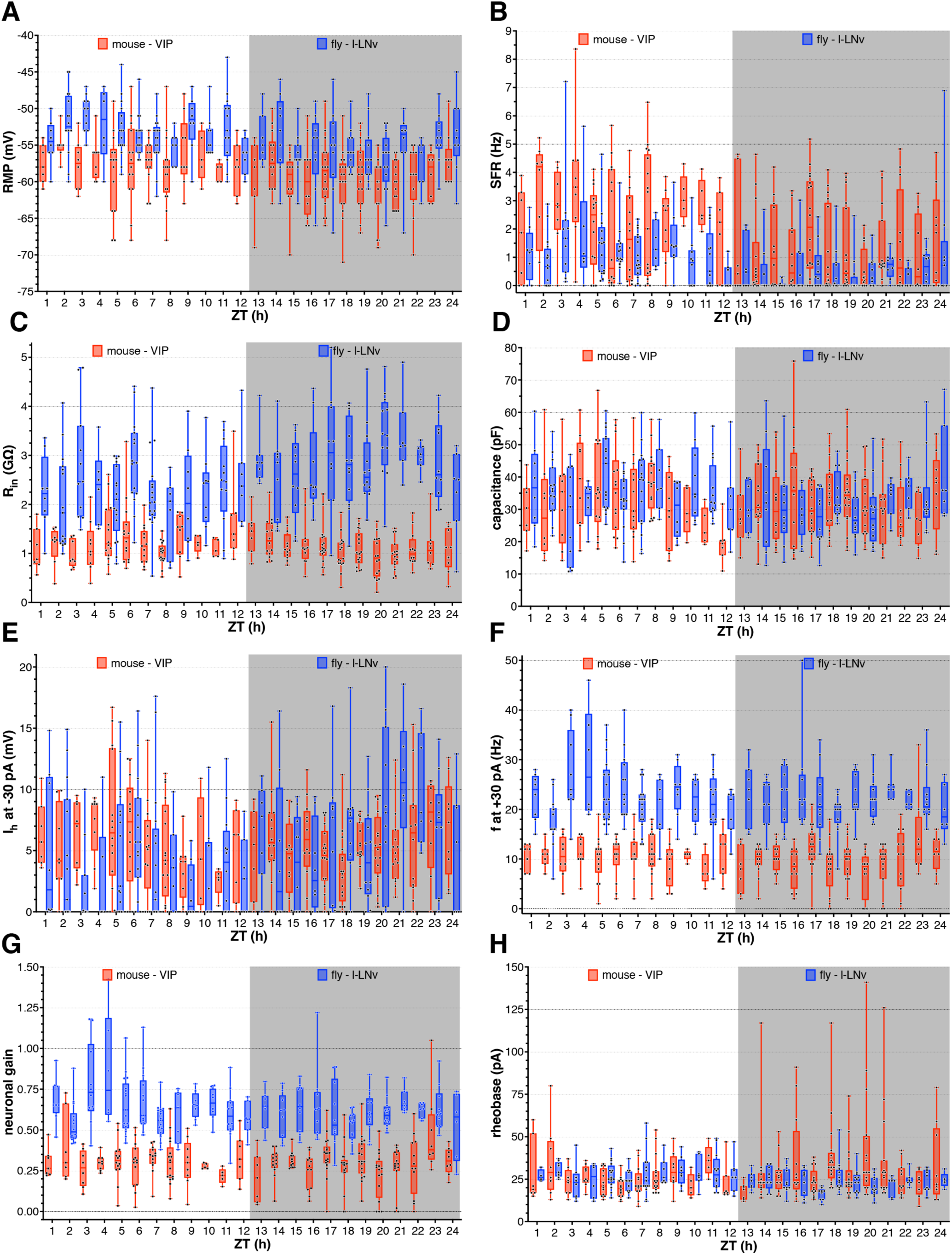
Individual data for diurnal variation in neurophysiological parameters of mouse and fly clock neurons. (**A**-**L**) Diurnal profiles of resting membrane potential (RMP, **A**), spontaneous firing rate (SFR, **B**), input resistance (R_in_, **C**), capacitance (**D**), h-current (I_h_ at -30 mV, **E**), firing rate at +30 pA (f at +30 pA, **F**), neuronal gain (**G**), rheobase (**H**) across 24 h under LD conditions. Box and whiskers, min-max, median indicated; n, see Table 1.

## Acknowledgements

We thank Dr Beatriz Bano-Otalora and Aadhithyan Babu for technical assistance. This work was supported by the BBSRC (BB/W000865/1, BB/Z517458/1, and BB/Z516594/1).

## Author contributions

Conceptualization: EB, HDP, KTA, MDCB, JJLH

Methodology: EB, MDCB

Validation: EB, MDCB

Formal analysis: EB

Investigation: EB

Data curation: EB

Resources: CN

Visualization: EB

Writing - original draft: EB

Writing - review & editing: EB, HDP, KTA, MDCB, JJLH

Supervision: HDP, KTA, MDCB, JJLH

Project administration: EB, JJLH

Funding acquisition: EB, HP, KTA, MDCB, JJLH

## Abbreviations

aCSF: artificial cerebrospinal fluid
AVP: arginine vasopressin
Cry: cryptochrome
I_A_: A-type potassium current
I_h_: hyperpolarisation-activated current
I_sus_: sustained potassium current
LD: light–dark
l-LNv: large ventrolateral neuron
LJP: liquid junction potential
MESOR: midline estimating statistic of rhythm
PDF: pigment dispersing factor
PIR: post-inhibitory rebound
R_in_: input resistance
RMP: resting membrane potential
SCN: suprachiasmatic nucleus
SFR: spontaneous firing rate
TTX: tetrodotoxin
VIP: vasoactive intestinal peptide
ZT: Zeitgeber time

